# The effect of sociality on competitive interactions among birds

**DOI:** 10.1101/2022.05.09.491173

**Authors:** Ilias Berberi, Eliot T. Miller, Roslyn Dakin

## Abstract

Sociality can provide many benefits, including increased foraging success, reproductive opportunities, and defence against predation. How does sociality influence the dominance hierarchies of ecological competitors? Here, we address this question using a large citizen science dataset of competitive interactions among birds foraging at backyard feeders, representing a network of over 55,000 interactions among 68 common species. We first show that species differ in average group size (the number of conspecifics observed together) as a fundamental measure of sociality. When analyzing heterospecific competition, we find that sociality is inversely related to dominance. On average, a single individual from a solitary species is more likely to displace a size-matched opponent than a single individual from a social species. Yet, we find that social species gain an increase in their competitive advantage when in the presence of their conspecifics, which may occur as a result of dynamics within their groups. Finally, we show that more social species have relatively fewer dominance interactions with heterospecifics, and more with conspecifics. Overall, these results demonstrate that sociality can influence competition in ecological networks. More social species have decreased competitive ability as individuals, but they may gain competitive ability in groups.

## INTRODUCTION

Animal sociality can provide many ecological and evolutionary advantages, including increased foraging success [1,2], more frequent mating opportunities [3,4], and improved survival through dilution and confusion effects on predators [5,6]. However, sociality may also incur costs under certain circumstance, such as conflict among conspecific group members that share a similar niche [7,8]. The causes and consequences of animal sociality have historically been of interest in evolutionary biology, with several different hypotheses for benefits that may explain the evolution of social grouping. One hypothesized benefit of sociality is that social coalitions may provide an advantage during conflict with competitors [9,10]. While both inter- and intraspecific dominance is often enhanced by body size [11,12] and age/experience [13], there may also be opportunities for subordinate individuals to gain an advantage over dominant competitors through social coalitions [9,14]. During foraging, animals face competition from both conspecifics and heterospecifics, but the influence of social grouping on the structure of these competitive systems has not yet been described. Our aim in this study is to evaluate how sociality influences competition in communities where interactions occur both within and between species.

One system that provides an opportunity to study this is the network of competitive interactions among birds at backyard bird feeders. In many temperate systems, survival in winter depends on acquiring adequate food resources [15,16]. Moreover, the use of artificial food sources (e.g., bird feeders) during winter foraging has been identified as having important evolutionary consequences in birds [17,18], including high rates of competitive interactions among and within species foraging at feeder locations [19,20]. These levels of competition may depend on the rate of resource depletion, where regularly depleted feeders may generate higher levels of resource competition than feeders that are continually replenished [21]. In the context of sociality, group foraging can provide more opportunities for resource acquisition than foraging independently due to shared vigilance [22,23] (but see [24]), social information gathering [25,26], and improved dominance over solitary foragers [27]. Individuals in many species have been found to adopt social foraging strategies in the winter when resources are scarce [28]. Additionally, sociality in the winter has been associated with reproductive advantages in subsequent mating seasons [29,30].

Here, we investigate how sociality influences competition using interactions among birds that visit backyard feeders. Our predictions at the outset of this study were that more social species may gain a benefit in interspecific conflict, at a cost of increased rates of intraspecific conflict. We tested these predictions using data from Project FeederWatch (PFW), a surveying program with a network of tens of thousands of observers across North America who monitor the presence and interactions of hundreds of bird species throughout winter (Figure 1). This project has yielded a rich dataset indicating who competes with whom, and how often (Figure 1D). Previous research using PFW observations has revealed that species body mass is the primary predictor of competitive dominance within winter bird communities across North America [12,31]. Owing to the large number of interactions observed and variation in competitive outcomes (Figure 1D), the PFW citizen science program provides an ideal system to test how variation in sociality might influence competitive interactions after accounting for the effects of body size.

**Figure 1.**
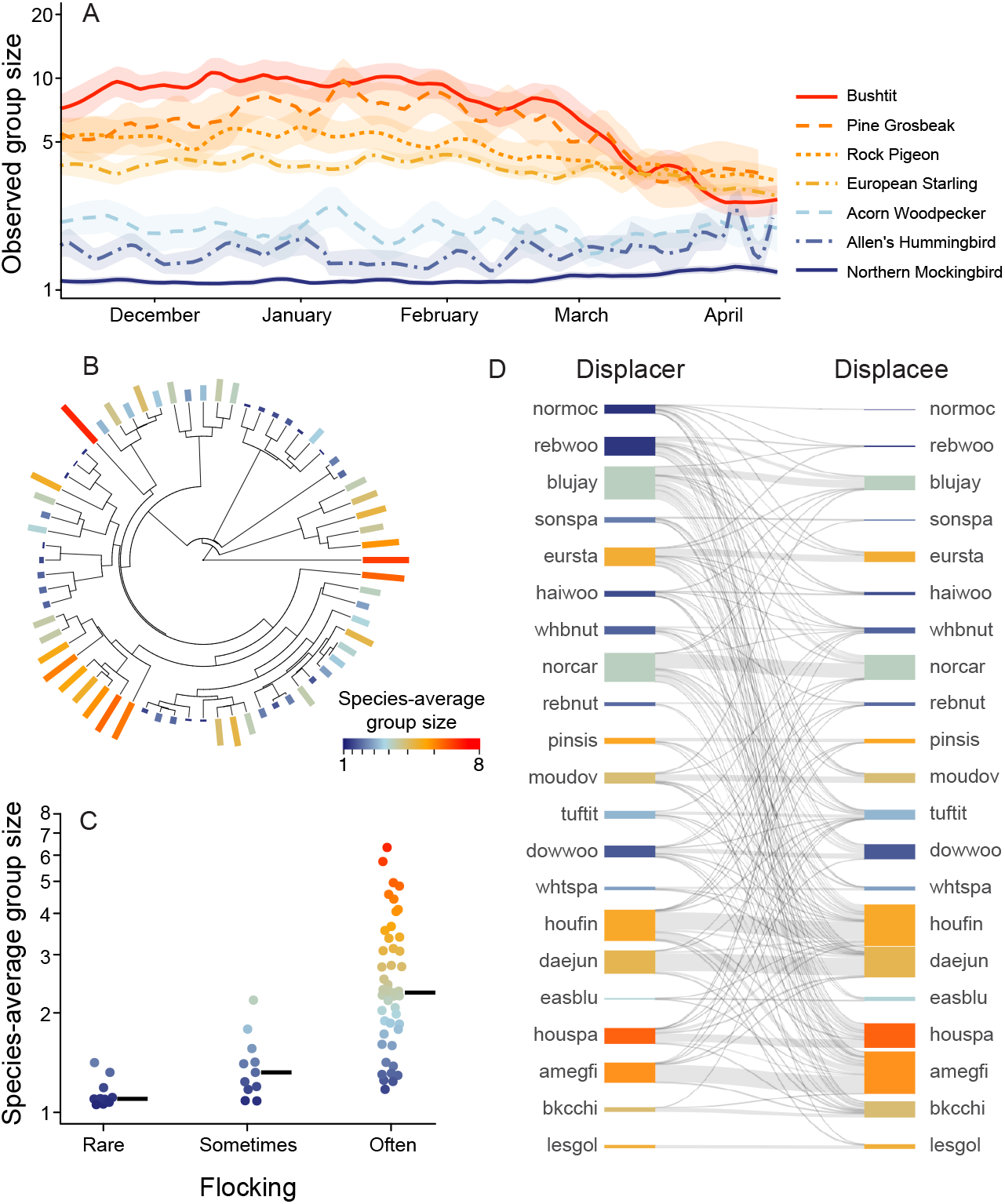
Bird species differ in winter sociality and competitive dominance. A) Example of observed group sizes (daily averages ± SE) for seven bird species during winter 2017-18, with species identity indicated by the line type. These species were chosen for visualization purposes to represent a wide range of avian families. B) Phylogeny of 68 focal bird species with species-average group size shown by the size and colour of the outer bars. Note that the colour gradient throughout this figure follows the legend shown in (B). C) Species-average group size in relation to descriptions from natural history accounts. Horizontal lines indicate medians. D) Competitive network illustrating dominance interactions among a subset of common species, chosen for visualization purposes, with at least 2,000 displacement interactions. Each row represents a species, and the thickness of grey connecting lines indicates the frequency of competitive interactions. Horizontal connections represent competitive interactions with conspecifics. Species nodes are ordered vertically (top to bottom) from the most to least dominant and coloured by average group size. Labels represent short-form common names (see Table S1 for details).

Here, we define species differences in winter sociality using a simple metric of conspecific group size (i.e., the number of individuals of a given bird species observed at a given location at the same time in the PFW database; Figure 1A-C) [32]. It is important to note that sociality also has other dimensions beyond the simple tempo-spatial aggregation studied here, such as the stability of group membership, the types of relationships between group members, and other emergent properties of group organization [33,34]. Nevertheless, conspecific groupings – individuals found in the same place at the same time – represent the most fundamental dimension of sociality that we expect to influence competitive outcomes across species.

As a first step in this study, we established that winter sociality differs across bird species by quantifying the repeatability of observed group sizes (Figure 1A). We also estimated the phylogenetic signal of species-average group size, which describes the extent to which this trait is more similar among closely related species (Figure 1B). Next, we analyzed the outcome of >55,000 heterospecific dominance interactions (a subset of these is shown in Figure 1D). Our aim was to evaluate whether average group size (as a species-level trait) is associated with species differences in dominance against size-matched opponents, and whether the presence of conspecifics would positively or negatively impact each species’ competitive ability (hereafter, we refer to this impact as the “conspecific effect”). We expected that more social species would have greater dominance over size-matched opponents, and a more positive conspecific effect. Finally, we analyzed the frequencies of conspecific and heterospecific interactions (Figure 1D), to determine whether sociality is associated with variation in the most common sources of conflict.

## MATERIALS AND METHODS

### (a) Data sources

We used a dataset collected by Project FeederWatch (PFW), an extensive, comprehensive and ongoing monitoring program with over 9 million observations of birds at winter feeding locations in North America during the months of November through April. For this study, we collected PFW data from the period November 2015 – April 2020. Participants were invited to monitor their chosen feeding location during weekly two-day observation periods throughout the winter. Participants recorded which species they observed and the maximum number of cooccurring individuals that they observed simultaneously at the feeding location for each species. Some participants also recorded displacement interactions, used in our analysis of competition described in section (c) of the Methods below. Additionally, as a measure of observation effort, each participant indicated the number of two-hour monitoring sessions performed (up to a maximum value of four monitoring sessions per two-day observation period). Collected PFW data routinely undergo a data validation protocol (highlighted in [35]) to reduce biases in species reporting and identification often associated with citizen science [36]. For the purpose of the present study, we focused our analyses on a set of the most common species that had at least 100 displacement interactions (n = 68 species, Table S2). All data analyses were performed in R (version 4.0.3) [37]. A detailed flow chart illustrating our main analysis methods is provided in the supplement (Figure S1).

### (b) Quantifying species differences in sociality

As our measure of sociality, we used group size, defined as the maximum number of individuals of a given species observed simultaneously during a given monitoring session (see examples in Figure 1A). To assess whether focal species differ in their average winter sociality, we analyzed the repeatability of observed group size values (e.g., Figure 1A), [38]defined as the proportion of total variation in observed group sizes that is attributed to differences among species. A high value of repeatability in this analysis indicates that most of the variation in group size is due to differences among species, whereas a low value (near 0) indicates that species differences in this trait are not detected. Because group sizes change with the onset of spring, we included dates from November 1 – February 28 (note that conclusions are unchanged when including a broader span of dates). Because the size of the dataset was so large (n > 6.1 million observations of the 68 focal species), we used a bootstrapping approach to estimate repeatability. For each bootstrap iteration, we randomly selected a subset of data with 1,000 observed group sizes for each of the 68 focal species (randomly chosen from the full dataset available for each species). We used this subset to compute conditional repeatability of observed group size using the “rptR” package [38], adjusted by observation effort as a fixed effect, and accounting for region (defined by latitude and longitude to the nearest degree), using the Poisson distribution and log link for count data. This procedure was repeated 100 times to derive a distribution of repeatability estimates for species differences in average group size.

Upon finding that observed group size differs among species, we computed estimates of species-average group size, which we use in further comparative analyses of our 68 focal species (see Figure S1 for details). To do this, we fit a regression model for each focal species where the response variable was the log-transformed observed group size (n = 2,257 – 414,471 observations, depending on the species), with a fixed effect of observation effort, and a random effect of region. From each fitted model, we defined species-average group size as the predicted group size given an effort of one monitoring session (Figure 1B; Table S2). We also estimated Pagel’s lambda for the 68 species-average group size values [39]. Lambda values close to 1 would indicate that closely related species have similar species-average group sizes, whereas lambda values near 0 would indicate that variation in species-average group size is independent of phylogenetic relatedness.

As an additional check on our metric of species-average group size, we collected information on the natural history each focal species during winter from species accounts in the Birds of the World online resource [40]. We used these descriptive accounts to classify each focal species into one of three categories: often flocking (clear descriptions of winter flocking and/or group behaviour), sometimes flocking (indicators of occasional winter flocking and/or participation as a satellite member), and rarely flocking (described as independent in winter).

### (c) Modelling heterospecific competition

We modelled heterospecific dominance interactions by using a dataset of displacement events, defined as instances when one individual purposefully takes over, or dislodges, another bird from the feeding location (see supplement for further details). PFW participants recorded 88,988 displacement interactions among 196 bird species in winters of 2016-17, 2017-18, 2018-19 and 2019-20, with most of these observations (71%) collected in 2017-18 and 2018-19. In this analysis, we used displacement interactions that involved two different species, wherein at least one of the two interactors belonged to a focal species (n = 55,483 interactions). It is important to note that these interactions involved one individual bird displacing another, and did not include collective events such as mobbing (see supplement for details).

We used a binomial mixed-effects regression model in the “lme4” package [41] to model the outcome of these heterospecific interactions. The aim of this model was to evaluate whether average group size (as a species-level trait) is associated with species differences in dominance against size-matched opponents, and whether the presence of conspecifics impacts competitive ability differently across species. The model had random effects for the identity of the “primary” species, and the identity of its opponent species. These identities were assigned randomly in a bootstrapping procedure, with details below. We included an additional random effect of region. The response variable was the outcome of the displacement interaction (i.e., whether the primary species was dominant in the displacement interaction). The model included fixed effects of the primary species’ average body mass (log-transformed), the primary species’ average group size (log-transformed), its relative group size during that particular observation session (expressed relative to its species average group size), and the relative body mass of its opponent (defined as the difference in log-transformed species-average body masses, opponent – primary). In this binomial model, the slope parameter for the primary species’ average group size captures the association between species-level differences in sociality and species-level differences in dominance (we also describe further comparative analyses of this relationship in section (d) below). The slope parameter for relative group size captures the “conspecific effect”. A positive estimate for this slope parameter would indicate that the presence of additional conspecifics can boost competitive ability, whereas a negative slope would indicate that the presence of conspecifics can hinder competitive ability. To test whether the conspecific effect is species-specific, we included a random slope for the relative group size parameter. This allows the conspecific effect to vary among species. We then compared the fit of the model with this random slope to a simpler model that lacked the random (species-specific) slope using AIC and the likelihood ratio test. Stronger support for the random slope model would indicate that the conspecific effect varies significantly among species.

To account for the fact that both species in a heterospecific interaction may be considered as primary, we bootstrapped this analysis by repeating the procedure described above 100 times. In each bootstrap iteration, the primary and opponent positions for each interaction were randomly assigned. Hence, the bootstrapping procedure ensured that each displacement interaction was only represented once, such that the analysis did not pseudoreplicate the data.

### (d) Comparative analyses of species dominance in relation to sociality

To test our main hypotheses about the relationships between dominance and winter sociality, we used the bootstrapped model fits to quantify two further species traits for comparative analysis: the ‘competitive dominance’ and ‘conspecific effect’ of each focal species. Competitive dominance was defined as the predicted probability that a single focal individual (group size of 1) would be the displacer in a dominance interaction against a same-sized (equal body mass) heterospecific opponent. We extracted these predictions from the 100 bootstrapped models described in Methods section (c) above, and took the average for each species as our measure of its competitive dominance when alone and facing size-matched opponents (see flow chart in Figure S1 for details).

To examine variation in the conspecific effect, we extracted the random (species-specific) slopes for relative group size. Negative slope values indicate that a species’ dominance decreases in the presence of conspecifics, whereas positive slope values indicate that a species’ dominance increases in the presence of conspecifics. A value of 0 would indicate that the presence of conspecifics does not affect a given species’ dominance. As above, we extracted these values from the 100 bootstrapped models in Methods section (c), and took the average for each species as its ‘conspecific effect’. We consider analyses of this trait to be conservative because the PFW observers reported only one group size per species per monitoring session (the maximum group size they observed). As a result, our estimates for the conspecific effect hinge on the assumption that species group sizes are relatively stable at a given feeder within a short span of time (e.g., [42–44]). Alternatively, if species group sizes are highly variable over the span of a few hours, it would reduce our ability to detect conspecific effects.

To test the relations between species dominance and sociality, we ran two comparative analyses using the “brms” package [45] (n = 68 focal species). The response variables were species’ competitive dominance, and the conspecific effect, respectively, modelled with Gaussian error distributions. The predictor of interest was species-average group size (log transformed). We ran these models both with and without accounting for the phylogenetic relationships among species in the random effects structure. If the phylogenetically adjusted model estimated a weaker effect size for a given predictor than the simpler model, it would indicate that the relationship is driven by differences among broader lineages. We ran models for 10,000 iterations using two chains and verified that Rhat values were all between 1.00 and 1.05, indicating model convergence. We verified that residuals for Gaussian regression models were normally distributed with no heteroscedasticity. To check whether our conclusions were robust to error in the y-values (Figure 2B,E) we repeated these comparative analyses 100 times using the full distribution of y-values (repeating the analyses once for each unique set of bootstrapped y-values). We found that the conclusions of the comparative analyses were consistent each time.

**Figure 2.**
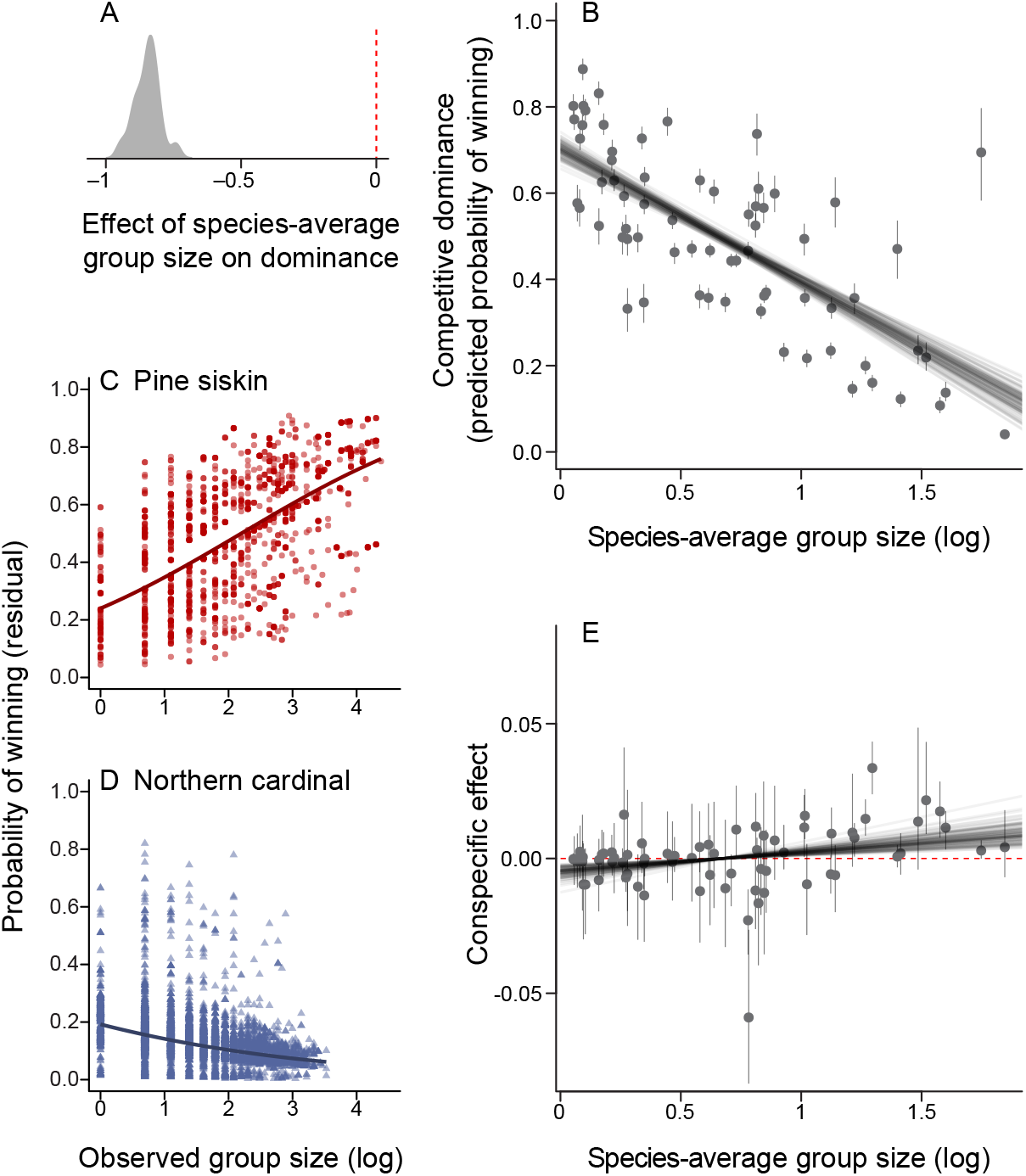
Relationship between sociality and heterospecific dominance. A) There is a strong negative association between species-average sociality and heterospecific dominance. This density plot shows the distribution of effect sizes from the bootstrapped models of heterospecific interactions in PFW. B) Scatterplot showing each species’ competitive dominance (the predicted probability of a single individual winning a heterospecific competition) in relation to species-average group size (n = 68 focal species). C-D) Examples illustrating how species differ in the relationship between dominance and the presence of conspecifics. Panels C-D show partial residual plots for two species: the pine siskin, where larger group sizes are associated with increased competitive ability, and the northern cardinal, where larger group sizes are associated with reduced competitive ability. E) Scatterplot showing the conspecific effect in relation to species-average group size (n = 68 focal species). In B and E, the y-values show bootstrapped averages (error bars represent 2.5^th^ and 97.5^th^ percentiles), and the grey lines show model fits based on 100 bootstrapped estimates (see methods for details).

### (e) Sources of conflict

Sociality may also influence the occurrence of competition (as well as its outcome). To compare the typical sources of conflict for each species with those expected by chance, we used a null model that was spatially constrained by region. The null model was computed as follows: within each *i^th^* region (defined by latitude and longitude rounded to the nearest degree), we first determined the proportion of total interactors for each *j^th^* focal species, *p_ij_*. Then, we calculated the expected number of conspecific and heterospecific interactions for each species, under the assumption that interaction rates are determined by the composition of interactors in that region. We took *p_ij_^2^* as the expected probability of a conspecific interaction, and 2 × *p_ij_* × (1 – *p_ij_*) as the expected probability of a heterospecific interaction for each species-region. We then multiplied these probabilities by the number of interactions in the region to obtain expected counts. Finally, we took the sum of the expected counts across all regions to get the total expected frequencies for each focal species and compared these null expectations with observed values.

To test whether a species’ social behaviour predicts variation in the sources of conflict, we fit a regression model where the response variable was the observed proportion of interactions against heterospecific opponents (one value per focal species). The predictor in this analysis was the species-average group size. This model was fit using the Bayesian brms package as described above, and we examined the model with and without accounting for the phylogenetic relationships among focal species. We verified that residuals for this model were normally distributed with no heteroscedasticity.

## RESULTS

### (a) Species differ in the degree of sociality

Observed group size varies both within and among species (Figure 1A), as indicated by significant repeatability (*R* _link-scale_ = 0.37, range 0.36-0.37, bootstrapped analysis of n = 68,000 observed group size values). Species-average group size has a very strong phylogenetic signal (lambda = 0.99, p < 0.001, n = 68, one value per focal species), indicating that closely related species share a very similar degree of sociality (Figure 1B). Species-average group size values also correspond to broad descriptions from natural history accounts (Figure 1C).

### (b) Species dominance in relation to sociality

In the bootstrapped model of all interactions, we found a strong negative association between heterospecific dominance and species-average sociality (Figure 2A). Consistent with this result, comparative analysis confirmed that competitive dominance is negatively related to sociality (Figure 2B, Table S3). Hence, a single individual from a more social species is much less likely to dominate size-matched opponents, as compared to a single individual from a less social species. This conclusion is upheld when accounting for phylogeny (Table S3).

We found strong support for species differences in the ‘conspecific effect’ (bootstrap average of ΔAIC = 40.2 in favour of random slope models; all LRT p-values < 0.006). Some species, like the pine siskin (*Carduelis pinus*), gain a large increase in competitive ability when more conspecifics are present (Figure 2C). Other species, like the northern cardinal (*Cardinalis cardinalis*), become less dominant when more conspecifics are present (Figure 2D). We found that the conspecific effect tends to be more positive in more social species (Figure 2E, Table S4). Thus, more social species gain a greater boost in dominance from the presence of conspecifics, as compared to less social species. This association, while weak, was upheld when accounting for phylogeny (Table S4).

### (c) Sources of conflict

Nearly all species experience less conflict with heterospecifics, and more conflict with conspecifics, than expected in a null model (Figure 3A-B). When examining species differences in the sources of conflict, we find that more social species have proportionately less conflict with heterospecifics (and hence, a greater proportion of their conflict is with conspecifics; Figure 3C, Table S5). This association was weaker when accounting for phylogeny, indicating that it is primarily driven by differences among lineages within our sample (Table S5).

**Figure 3.**
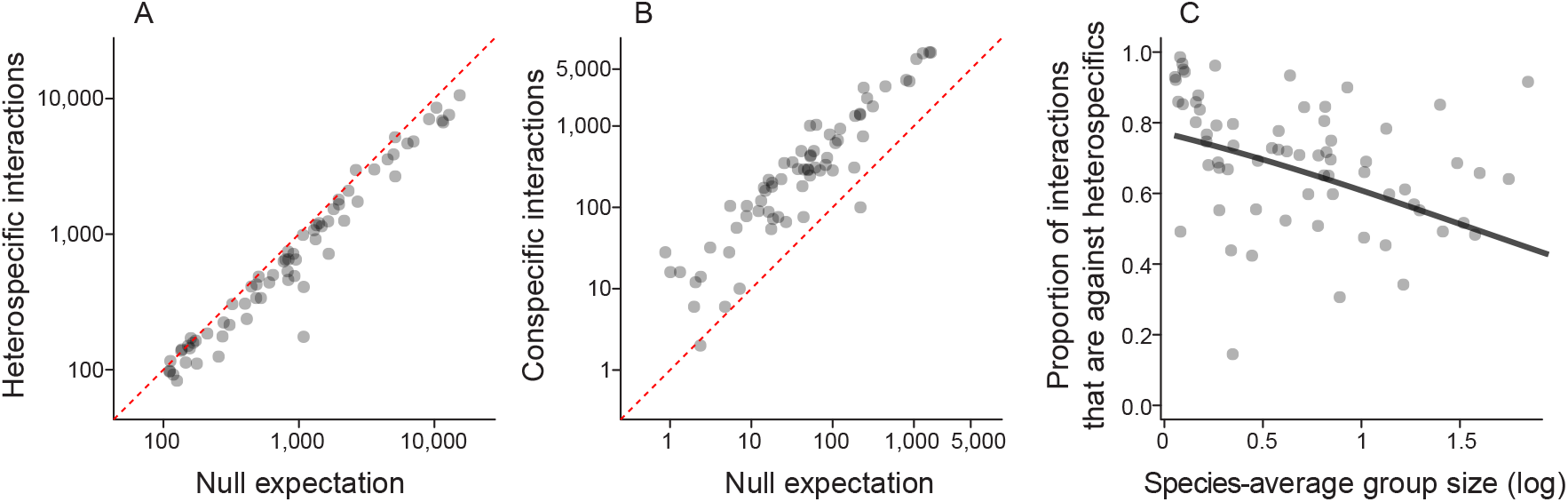
Conflict with heterospecifics occurs less often than expected, especially in more social lineages. Observed frequencies of (A) heterospecific and (B) conspecific dominance interactions in relation to null expected values for 68 focal species. Along the red dotted 1:1 lines, the observed frequency equals the expected value. (C) Scatterplot showing the proportion of conflict with heterospecifics in relation to species-average group size. Note that the line of best fit does not account for phylogeny (see Table S5 for additional details).

## DISCUSSION

Here, we examine the effect of social clustering on competition among winter resident birds. We find that species vary in group sizes, and that certain phylogenetic clades are more social than others. Contrary to our initial prediction, we find that individuals from more social species are also weaker competitors when facing size-matched heterospecific opponents (Figure 2B). This result suggests that there may be an evolutionary trade-off whereby the evolution of greater sociality is associated with reduced competitive ability of an individual against other species [31]. Our results also show that more social species tend to experience an increase in competitive ability when in the presence of their conspecifics (Figure 2E). Collectively, these results suggest that more social species have reduced competitive ability as individuals, but gain a boost in competitive ability in groups.

We can hypothesize several potential mechanisms that may drive the patterns in Figure 2. First, the strong decrease in competitive dominance observed in more social species (Figure 2B) may be due to a loss of aggressive or antagonistic behaviours against heterospecifics during the evolution of sociality. It is also possible that social species simply invest less into competition with other species over food, for example if it is more beneficial for social species to limit their interactions to conspecific competitors, or if social species are less motivated to compete because they can rely on other strategies for finding food (e.g., social information transfer) [46,47]. To test these hypotheses, further studies are required to test behavioural differences under conditions where motivation and experience can be controlled.

We can also propose several hypotheses that may explain variation in the conspecific effect, and the fact that it is generally more positive in more social species (Figure 2E). One possibility is that the conspecific effect may be driven intrinsically, as a consequence of other species-level traits. For example, social species may have increased plasticity in aggression, boldness, and/or vigilance that allows them to adjust their agonistic behaviour depending on the context [48,49]. The evolution of sociality has also been shown to be associated with enhanced cognition [50] and collective intelligence [51], which could also underly this effect. Yet another possibility is that the conspecific effect could be a by-product of within-species dominance hierarchies in more social species. For example, when a social flock visits a feeder, access may be biased toward the most dominant and aggressive individual(s) in the group, who may in turn be more likely to prevail against heterospecific competitors. The conspecific effect may also be driven extrinsically by opponent responses (e.g., if larger groups of a more social species are more coordinated in ways that generate more intimidating visual and/or acoustic signals). These intrinsic and extrinsic explanations for the conspecific effect are not mutually exclusive. Testing causal hypotheses will require careful experimentation.

Our results examining sources of conflict suggest that the evolution of sociality leads to less frequent conflict with heterospecific competitors, and more frequent conflict among conspecifics (Figure 3). In the presence of many conspecifics, individuals may interact more often, especially if they are collecting similar resources [7,8]. Members of a group may experience increased rates of aggressive interactions at times of food scarcity [52,53]. One way to reduce competition pressures between conspecifics may be through participation in mixed-species flocks. Mixed-species flocks have been found to provide members with greater advantages than conspecific coalitions, including increased foraging rates and reduce vigilance effort [1]. While some evidence suggests that members of mixed-species flocks are more similar than expected [54], they undoubtedly vary more in their ecological needs than same-species flocks and therefore may demonstrate a reduced level of within-group conflict than a conspecific coalition [55–57]. A key question for future research is whether mixed-species flocking can reduce the frequency of within-group conflict.

Our results raise interesting questions for future research on other ecological factors. For example, low resource availability is expected to generate higher levels of competition [21]. It would be interesting to test whether environmental harshness influences the magnitude of the conspecific effect. It is possible that social species experience even stronger conspecific effects when resources are more limited. Additionally, weather conditions such as precipitation and temperature may constrain foraging behaviour and may affect the costs and benefits of competitive interactions as well as social clustering behaviour [58]. Further studies are thus required to investigate how these environmental challenges may affect the structure of competitive dominance networks.

It is important to note that while we define sociality as the aggregation of individuals in space and time, there are other dimensions of sociality that are also expected to play a role in influencing competition. For example, members of a temporary flock with short-term social bonds may experience a weaker influence of conspecifics than members in a structured dominance hierarchy with long-term, persistent bonds [12,59–61]. This raises interesting questions for future work that can be addressed by tracking specific individuals within groups. For example, how does the strength of social bonds and the organization of a group influence a species’ competitive ability? Do such effects on competition change with temporal variation in species social organization? Within a species, how does the strength of a coalitionary relationship affect the dominance of one group over others?

Previous studies have examined the benefits of coalitionary behaviour on within-species conflict [9,14,62]. Our results here demonstrate how sociality can also influence the dynamics of ecological competition, and how a species’ tendency to form social groups can be associated with the evolution of reduced competitive ability as individuals. These findings provide valuable insight into the fundamental effects of grouping behaviour on competitive interactions during foraging. Further questions remain as to the effects of specific social bonds and the emergent properties of groups on competitive interactions, along with the effects of additional influential variables, such as cognitive ability, resource abundance, and environmental conditions on the relationships between sociality and competition.

## DATA ACCESSIBILITY

All data and R scripts are available at: https://figshare.com/s/1cd6708a27a541029409

The repository will be made public when the final version of the study is published.

## AUTHOR CONTRIBUTIONS

IB and RD designed the study with input from ETM. ETM developed the behavioural interactions methodology in Project FeederWatch. IB and RD analyzed the data and wrote the manuscript. All authors edited the manuscript.

## ACKNOWLEDGEMENTS

We would like to thank the Cornell Lab of Ornithology and Birds Canada for developing and managing Project FeederWatch and sharing the results. We especially thank the thousands of volunteers who have contributed their observations to Project FeederWatch, and we thank the peer reviewers for helpful feedback on this study.

