## supplement for "The effect of sociality on competitive interactions among birds"

### Definition of displacement events in Project FeederWatch

*"Displacement is when one bird (the "source") tries to take over a resource (usually food, but sometimes a perch) occupied by another bird (the "target"). A displacement event is successful if the source bird dislodges the target bird from a perch or feeder. The source bird needs to be purposefully attempting to take the perch of the target bird, rather than landing on a spot as the target bird was about to leave on its own accord. Displacement behavior does not include when a bird flies away to escape a predator or when a group of birds mobs another bird. Sometimes large birds, such as Blue Jays or Red-bellied Woodpeckers, can arrive suddenly at a feeder and cause other birds to scatter. Or sometimes a flock of birds, such as Bushtits, can arrive and cause other birds to leave. Such instances are difficult to interpret so we ask that you only report clear examples of one individual attempting to displace another individual."*

Project FeederWatch Detailed Instructions, Cornell Lab of Ornithology, https://feederwatch.org/about/detailed-instructions/#record-behavior-interactions

**Table S1.** Key for the species labels presented in Figure 1D.

| Label | Common name | Scientific name |
|---|---|---|
| normoc | Northern Mockingbird | *Mimus polyglottos* |
| rebwoo | Red-bellied Woodpecker | *Melanerpes carolinus* |
| blujay | Blue Jay | *Cyanocitta cristata* |
| sonspa | Song Sparrow | *Melospiza melodia* |
| eursta | European Starling | *Sturnus vulgaris* |
| haiwoo | Hairy Woodpecker | *Picoides villosus* |
| whbnut | White-breasted Nuthatch | *Sitta carolinensis* |
| norcar | Northern Cardinal | *Cardinalis cardinalis* |
| rebnut | Red-breasted Nuthatch | *Sitta canadensis* |
| pinsis | Pine Siskin | *Carduelis pinus* |
| moudov | Mourning Dove | *Zenaida macroura* |
| tuftit | Tufted Titmouse | *Baeolophus bicolor* |
| dowwoo | Downy Woodpecker | *Picoides pubescens* |
| whtspa | White-throated Sparrow | *Zonotrichia albicollis* |
| houfin | House Finch | *Carpodacus mexicanus* |
| daejun | Dark-eyed Junco | *Junco hyemalis* |
| easblu | Eastern Bluebird | *Sialia sialis* |
| houspa | House Sparrow | *Passer domesticus* |
| amegfi | American Goldfinch | *Carduelis tristis* |
| bkcchi | Black-capped Chickadee | *Parus atricapillus* |
| lesgol | Lesser Goldfinch | *Carduelis psaltria* |

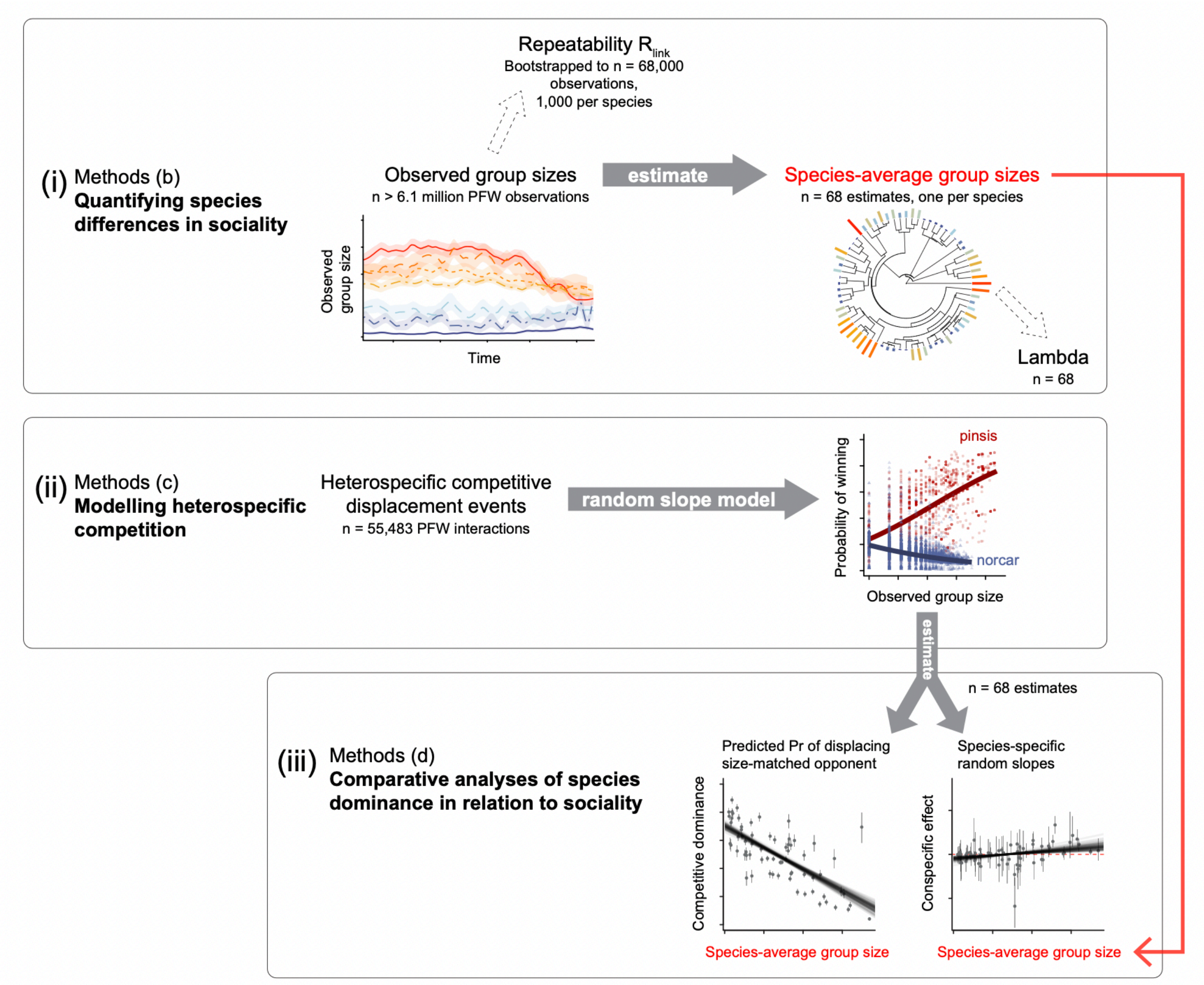


**Figure S1.** Flow chart illustrating the methods used to (i) quantify species differences in sociality, (ii) model heterospecific competition, and (iii) conduct a comparative analysis of the relationship between species-average competition traits and species-average group size. The labels (b), (c) and (d) refer to the relevant sections of the main text Methods. Figure panels within each row are taken from the main text, to help illustrate the workflow. Note that for clarity, only a subset of the focal species are shown in figures on the top left (taken from main Figure 1A) and the middle row (taken from main Figure 2C-D).

**Table S2.** Descriptive statistics of group sizes for the 68 focal species in the analyses, from November–February inclusive. Observed group sizes are calculated from the raw data. Species-average group sizes are estimated from models as described in the main text Methods.

| Common name | Scientific name | Observed group size (mean) | ± SD | Percentile 25th | Percentile 75th | Species-average group size |
|---|---|---|---|---|---|---|
| Acorn Woodpecker | *Melanerpes formicivorus* | 2.2 | 1.3 | 1 | 3 | 1.9 |
| Allen's Hummingbird | *Selasphorus sasin* | 1.8 | 1.9 | 1 | 2 | 1.3 |
| American Crow | *Corvus brachyrhynchos* | 3.4 | 5.5 | 1 | 4 | 2.3 |
| American Goldfinch | *Spinus tristis* | 7.4 | 10.3 | 2 | 8 | 4.1 |
| American Robin | *Turdus migratorius* | 5.1 | 14.4 | 1 | 5 | 2.2 |
| American Tree Sparrow | *Spizelloides arborea* | 3.8 | 4.9 | 1 | 4 | 2.3 |
| Anna's Hummingbird | *Calypte anna* | 2.1 | 3.0 | 1 | 2 | 1.4 |
| Baltimore Oriole | *Icterus galbula* | 2.8 | 3.0 | 1 | 4 | 1.1 |
| Band-tailed Pigeon | *Patagioenas fasciata* | 8.0 | 10.3 | 2 | 10 | 3.1 |
| Bewick's Wren | *Thryomanes bewickii* | 1.1 | 0.4 | 1 | 1 | 1.1 |
| Black-billed Magpie | *Pica hudsonia* | 2.9 | 2.8 | 1 | 3 | 1.6 |
| Black-capped Chickadee | *Poecile atricapillus* | 3.4 | 3.2 | 2 | 4 | 2.8 |
| Blue Jay | *Cyanocitta cristata* | 3.3 | 3.1 | 1 | 4 | 2.2 |
| Brown-headed Cowbird | *Molothrus ater* | 8.5 | 19.6 | 1 | 8 | 2.8 |
| Brown-headed Nuthatch | *Sitta pusilla* | 1.5 | 0.7 | 1 | 2 | 1.3 |
| Brown Thrasher | *Toxostoma rufum* | 1.1 | 0.3 | 1 | 1 | 1.1 |
| Bushtit | *Psaltriparus minimus* | 12.1 | 7.5 | 6 | 16 | 6.3 |
| California Scrub-Jay | *Aphelocoma californica* | 2.1 | 1.8 | 1 | 2 | 1.8 |

| | | | | | | |
|---|---|---|---|---|---|---|
| Carolina Chickadee | *Poecile carolinensis* | 2.1 | 1.2 | 1 | 3 | 1.8 |
| Carolina Wren | *Thryothorus ludovicianus* | 1.3 | 0.6 | 1 | 2 | 1.2 |
| Cassin's Finch | *Haemorhous cassinii* | 4.8 | 5.8 | 1 | 6 | 2.4 |
| Chestnut-backed Chickadee | *Poecile rufescens* | 2.9 | 2.1 | 1 | 4 | 2.5 |
| Chipping Sparrow | *Spizella passerina* | 6.7 | 10.9 | 1 | 8 | 2.0 |
| Common Grackle | *Quiscalus quiscula* | 12.4 | 74.5 | 1 | 7 | 2.3 |
| Common Redpoll | *Acanthis flammea* | 16.4 | 23.3 | 2 | 21 | 4.8 |
| Curve-billed Thrasher | *Toxostoma curvirostre* | 1.7 | 2.2 | 1 | 2 | 1.1 |
| Dark-eyed Junco | *Junco hyemalis* | 6.8 | 7.3 | 2 | 8 | 3.1 |
| Downy Woodpecker | *Dryobates pubescens* | 1.7 | 0.9 | 1 | 2 | 1.3 |
| Eastern Bluebird | *Sialia sialis* | 2.9 | 2.7 | 2 | 4 | 2.0 |
| Eastern Towhee | *Pipilo erythrophthalmus* | 1.6 | 0.9 | 1 | 2 | 1.2 |
| Eurasian Collared-Dove | *Streptopelia decaocto* | 4.6 | 6.1 | 2 | 5 | 2.4 |
| European Starling | *Sturnus vulgaris* | 8.6 | 26.3 | 2 | 8 | 3.4 |
| Evening Grosbeak | *Coccothraustes vespertinus* | 14.1 | 17.7 | 3 | 19 | 4.6 |
| Fox Sparrow | *Passerella iliaca* | 1.7 | 1.5 | 1 | 2 | 1.2 |
| Golden-crowned Sparrow | *Zonotrichia atricapilla* | 4.1 | 3.7 | 2 | 5 | 2.1 |
| Hairy Woodpecker | *Dryobates villosus* | 1.4 | 0.7 | 1 | 2 | 1.2 |
| House Finch | *Haemorhous mexicanus* | 5.6 | 6.4 | 2 | 7 | 3.6 |

| | | | | | | |
|---|---|---|---|---|---|---|
| House Sparrow | *Passer domesticus* | 10.7 | 13.1 | 3 | 14 | 5.0 |
| Lesser Goldfinch | *Spinus psaltria* | 7.1 | 7.7 | 2 | 9 | 3.4 |
| Mountain Chickadee | *Poecile gambeli* | 2.4 | 2.2 | 1 | 3 | 1.8 |
| Mourning Dove | *Zenaida macroura* | 6.4 | 7.4 | 2 | 8 | 2.8 |
| Northern Cardinal | *Cardinalis cardinalis* | 3.7 | 3.8 | 2 | 4 | 2.2 |
| Northern Flicker | *Colaptes auratus* | 1.5 | 1.0 | 1 | 2 | 1.2 |
| Northern Mockingbird | *Mimus polyglottos* | 1.1 | 0.5 | 1 | 1 | 1.1 |
| Orange-crowned Warbler | *Leiothlypis celata* | 1.2 | 0.6 | 1 | 1 | 1.1 |
| Painted Bunting | *Passerina ciris* | 3.4 | 2.6 | 1 | 5 | 1.6 |
| Pileated Woodpecker | *Dryocopus pileatus* | 1.2 | 0.5 | 1 | 1 | 1.1 |
| Pine Grosbeak | *Pinicola enucleator* | 9.3 | 8.9 | 3 | 12 | 4.4 |
| Pine Siskin | *Spinus pinus* | 9.1 | 16.0 | 2 | 10 | 3.7 |
| Pine Warbler | *Setophaga pinus* | 1.7 | 1.6 | 1 | 2 | 1.3 |
| Purple Finch | *Haemorhous purpureus* | 3.4 | 4.3 | 1 | 4 | 2.3 |
| Red-bellied Woodpecker | *Melanerpes carolinus* | 1.2 | 0.5 | 1 | 1 | 1.1 |
| Red-breasted Nuthatch | *Sitta canadensis* | 1.5 | 0.8 | 1 | 2 | 1.3 |
| Red-winged Blackbird | *Agelaius phoeniceus* | 11.0 | 24.2 | 1 | 10 | 3.1 |
| Rock Pigeon | *Columba livia* | 8.3 | 9.6 | 2 | 10 | 4.1 |
| Song Sparrow | *Melospiza melodia* | 1.6 | 1.3 | 1 | 2 | 1.4 |
| Spotted Towhee | *Pipilo maculatus* | 2.2 | 1.6 | 1 | 3 | 1.4 |
| Steller's Jay | *Cyanocitta stelleri* | 3.3 | 3.6 | 1 | 4 | 2.3 |

| | | | | | | |
|---|---|---|---|---|---|---|
| Townsend's Warbler | *Setophaga townsendi* | 1.3 | 0.8 | 1 | 1 | 1.2 |
| Tufted Titmouse | *Baeolophus bicolor* | 2.3 | 1.4 | 1 | 3 | 1.7 |
| Varied Thrush | *Ixoreus naevius* | 2.1 | 2.9 | 1 | 2 | 1.4 |
| White-breasted Nuthatch | *Sitta carolinensis* | 1.5 | 0.7 | 1 | 2 | 1.3 |
| White-crowned Sparrow | *Zonotrichia leucophrys* | 5.1 | 6.7 | 1 | 6 | 1.9 |
| White-throated Sparrow | *Zonotrichia albicollis* | 3.7 | 4.4 | 1 | 4 | 1.6 |
| White-winged Dove | *Zenaida asiatica* | 7.8 | 9.8 | 2 | 10 | 2.3 |
| Wild Turkey | *Meleagris gallopavo* | 10.6 | 10.1 | 4 | 14 | 5.7 |
| Yellow-bellied Sapsucker | *Sphyrapicus varius* | 1.1 | 0.3 | 1 | 1 | 1.1 |
| Yellow-rumped Warbler | *Setophaga coronata* | 1.8 | 2.1 | 1 | 2 | 1.4 |

**Table S3.** Comparative analysis of species differences in competitive dominance (n = 68 focal species). Two Bayesian regressions were fit, one with a random effect of phylogeny, and one without. Each model was run for 10,000 iterations.

| | Model without phylogeny | | | Model with phylogeny | | |
|---|---|---|---|---|---|---|
| | | 95% CI | | | 95% CI | |
| | Estimate | Lower | Upper | Estimate | Lower | Upper |
| Fixed effect | | | | | | |
| Intercept | 0.702 | 0.644 | 0.762 | 0.863 | 0.689 | 1.022 |
| Group size$_{[log]}$ | **-0.307** | **-0.377** | **-0.237** | **-0.347** | **-0.398** | **-0.291** |
| Random effect | - | - | - | - | - | - |
| Phylogeny | - | - | - | **0.017** | **0.014** | **0.020** |

**Table S4.** Comparative analysis of species differences in the conspecific effect (n = 68 focal species). Two Bayesian regressions were fit, one with a random effect of phylogeny, and one without. Each model was run for 10,000 iterations.

| | Model without phylogeny | | | Model with phylogeny | | |
|---|---|---|---|---|---|---|
| | | 95% CI | | | 95% CI | |
| | Estimate | Lower | Upper | Estimate | Lower | Upper |
| Fixed effect | | | | | | |
| Intercept | -0.005 | -0.010 | 0.000 | -0.006 | -0.014 | 0.002 |
| Group size$_{[log]}$ | **0.007** | **0.002** | **0.013** | **0.008** | **0.001** | **0.014** |
| Random effect | - | - | - | - | - | - |
| Phylogeny | - | - | - | **0.000** | **0.000** | **0.001** |

**Table S5.** Comparative analysis of species differences in the typical sources of conflict, analyzed as the proportion of interactions that were against heterospecific conspecific opponents (n = 68 focal species). Two Bayesian linear regressions were fit, one with a random effect of the phylogeny, and one without. Each model was run for 10,000 iterations.

| | Linear regression | | | Phylogenetic regression | | |
|---|---|---|---|---|---|---|
| | | 95% CI | | | 95% CI | |
| | Estimate | Lower | Upper | Estimate | Lower | Upper |
| Fixed effect | | | | | | |
| Intercept | 0.780 | 0.711 | 0.852 | 0.714 | 0.505 | 0.899 |
| Group size$_{[log]}$ | **-0.125** | **-0.211** | **-0.041** | -0.070 | -0.172 | 0.035 |
| Random effect | - | - | - | - | - | - |
| Phylogeny | - | - | - | **0.017** | **0.006** | **0.028** |

**Analyzing heterospecific competition using species-specific models**
In an earlier version of this study, we quantified each focal species' competitive dominance and conspecific effects using a different analytical pipeline that involved fitting 68 species-specific models (one for each focal species). Each model analyzed the outcome of heterospecific competitive interactions for a single focal species at a time. We then used these species-specific models to extract estimates of each focal species' competitive dominance and conspecific effect. Full details of this approach can be found in an earlier version of our study, available here: https://doi.org/10.1101/2022.05.09.491173

Following reviewer feedback, we have replaced this early version of the analysis with an improved approach that uses bootstrapping to analyze all heterospecific interactions at once, in a single large model. This model includes a random slope term to evaluate variation in the conspecific effect across species. With both approaches, we found the same conclusions. Hence, the conclusions of this study are robust to the choice of analytical pipeline used.
